# cAMP promotes acute lysosome biogenesis through TFEB nuclear import-export dynamics

**DOI:** 10.1101/2025.11.24.690233

**Authors:** Saumya Bhatt, Mohammad Ali Abbas Zaidi, Nicholas Woods, Cody N. Rozeveld, Micah B. Schott

## Abstract

Cyclic adenosine monophosphate (cAMP) signaling is a major stimulus for lipid and glucose catabolism, yet catabolic processes like these can also coordinate with lysosome-dependent degradation. However, the impact of cAMP signaling on lysosomal dynamics remains unclear. Transcription factor EB (TFEB), a master regulator of lysosomal biogenesis, is regulated by stimulus-dependent nuclear–cytoplasmic shuttling through a variety of phosphorylation events. Here, we find that elevating intracellular cAMP with forskolin and IBMX induces rapid nuclear import of TFEB-GFP within 30 minutes and coincides with a transient upregulation of TFEB target lysosome genes. By 8 hours, TFEB returns to the cytoplasm, accompanied by transcriptional downregulation. Inhibition of cAMP-dependent protein kinase A (PKA) using H89 did not block nuclear import but unexpectedly caused sustained nuclear accumulation, indicating that PKA promotes TFEB nuclear export. Consistent with this, phosphoproteomic profiling revealed increased phosphorylation of a PKA-consensus motif (RRx<u>S)</u> during the export phase. These findings suggest that cAMP–PKA signaling plays a novel role in temporally “tuning” lysosomal gene expression by regulating TFEB nuclear-cytoplasmic shuttling.

**Summary:** This study reveals that cAMP signaling dynamically regulates TFEB subcellular localization, promoting transient, calcium-dependent nuclear import as well as downstream, PKA-dependent export and phosphorylation at serines 466/467. These findings uncover a novel mechanism by which cAMP stimulation fine-tunes lysosomal gene expression by regulating TFEB nuclear import and export.

## Introduction

cAMP is a central second messenger that integrates hormonal and metabolic cues to regulate diverse processes, including lipid and glycogen metabolism(Ravnskjaer et al., 2016). Beyond these classic catabolic roles, cAMP signaling has also been implicated in lysosomal regulation, including lysosomal acidification and functional control(Li et al., 2019a; Rahman et al., 2016; Rodriguez et al., 1999). However, the mechanisms linking cAMP signaling to lysosomal dynamics remain poorly understood. Transcription factor EB (TFEB) is a master regulator of lysosome biogenesis and lipid droplet (LD) breakdown, while also regulating a range of vital functions, including autophagy, metabolism, stress response, cellular differentiation, and immune regulation(Franco-Juarez et al., 2022; Tan et al., 2022). Dysregulation of TFEB has been implicated in a broad spectrum of human diseases, including neurodegenerative disorders such as Alzheimer’s and Parkinson’s disease, cancer, and metabolic disorders such as liver steatosis, obesity, and diabetes(Cheng et al., 2023; Cortes and La Spada, 2019; Gebrie, 2023; Kao et al., 2022; Rai et al., 2021; Yan, 2022; Yang et al., 2023).

TFEB transcriptional activity is determined by its nucleocytoplasmic shuttling(Puertollano et al., 2018). Its subcellular localization is primarily controlled by post-translational modifications, with kinases such as mTOR, ERK, AKT, GSK3β, and PKC that modulate its phosphorylation state to determine nuclear versus cytoplasmic localization(Takla et al., 2023). For example, TFEB is phosphorylated at S211 by mTORC1 to sequester TFEB in the cytoplasm, inhibiting transcription of genes related to lysosome biogenesis and autophagy during anabolic growth. Dephosphorylation of S211, orchestrated by Ca^2+^/calcineurin phosphatase, triggers rapid import of TFEB into the nucleus(Zhao et al., 2024). Within the nucleus, TFEB then binds to Coordinated Lysosomal Expression And Regulation (CLEAR) elements present in the promoter of target genes that regulate autophagy and lysosomal biogenesis(Sardiello et al., 2009; Settembre et al., 2011). The duration of TFEB’s nuclear residence is critical for its transcriptional activity, as recent studies now show that TFEB nuclear export can also act as the key factor limiting this residence time, thereby “tuning” TFEB-driven gene expression(Napolitano et al., 2018; Ruolo et al., 2025). Indeed, dysregulation of the TFEB import-export cycle can lead to impaired gene activation(Park et al., 2023) or constitutive TFEB activity(Alesi et al., 2024). Interestingly, phosphorylation events not only mediate TFEB nuclear import, but can also drive nuclear export. For example, the nuclear export protein CRM1 (also known as Exportin 1) was recently shown to recognize an “inducible” nuclear export signal (NES) in TFEB’s N-terminal region that is unveiled by the hierarchical phosphorylation of TFEB by mTOR at S142 and GSK3β at S138(Li et al., 2018; Napolitano et al., 2018).

In addition to its N-terminal regulatory sites, TFEB’s C-terminus contains multiple phosphorylation sites that modulate its localization, stability, and transcriptional activity. For example, phosphorylation of S467 by AKT retains TFEB in the cytoplasm(Palmieri et al., 2017), whereas PKCβ-mediated phosphorylation at S461/S462, S466, and S468 stabilizes TFEB from proteasomal degradation(Ferron et al., 2013). However, AMPK phosphorylation of S466, S467, and S469 enhances TFEB’s transcriptional activity(Paquette et al., 2021). These findings collectively underscore the importance of S466 and S467 as convergence points for multiple signaling pathways that may exert different effects on TFEB subcellular localization and transcriptional activity.

In the current study, our sequence analysis revealed a consensus phosphorylation motif for cAMP-dependent protein kinase A (PKA) phosphorylation at S467. cAMP stimulation using Fsk and IBMX promotes rapid TFEB nuclear import within 30 minutes, leading to rapid induction of CLEAR genes, followed by persistent export back to the cytoplasm within 8 hours, coinciding with a reduction in transcription of TEFB target genes. Unexpectedly, the inhibition of PKA did not prevent cAMP-induced nuclear import but specifically blocked subsequent nuclear export. Phosphoproteomic and site-directed mutagenesis revealed phosphorylation of S466 and S467 as key residues that promote PKA-dependent nuclear export of TFEB following nuclear import. These findings define a cAMP–PKA–TFEB signaling axis that coordinates the temporal regulation of TFEB transcriptional activity and lysosomal dynamics.

## Results

### cAMP promotes TFEB nuclear import via calcium-signaling and increases lysosomal volume

To determine the impact of cAMP on TFEB localization and activity, HeLa cells stably expressing TFEB-GFP were treated with the adenylyl cyclase activator forskolin (Fsk, 10 μM) and the phosphodiesterase inhibitor 3-isobutyl-1-methylxanthine (IBMX, 0.5 mM) to elevate intracellular cAMP. Live-cell time lapse fluorescence microscopy revealed a rapid translocation of TFEB-GFP from the cytoplasm to the nucleus upon Fsk+IBMX treatment, with a substantial nuclear accumulation seen as early as 30 minutes (Fig. 1A). Quantification of nuclear-to-cytoplasmic TFEB fluorescence confirmed a significant increase in nuclear TFEB within 60 minutes compared with DMSO controls (Fig. 1B). A similar response was observed in Fsk+IBMX treated AML-12 hepatocytes transiently expressing TFEB-GFP, indicating that this response may also occur in hepatocytes (Fig. S1A-B). To assess whether TFEB nuclear localization correlates with lysosomal remodeling, HeLa-TFEB-GFP cells were labeled with LysoTracker Deep Red and imaged by live-cell confocal microscopy. cAMP elevation resulted in visibly increased lysosomal signal intensity (Fig. 1C). Quantification showed a significant increase in total lysosomal volume after 2 hours of treatment, while lysosome number remained unchanged (Fig. 1D-E). These findings suggest that TFEB translocation into the nucleus may correlate with TFEB transcriptional activity after cAMP stimulation.

**Figure 1:**
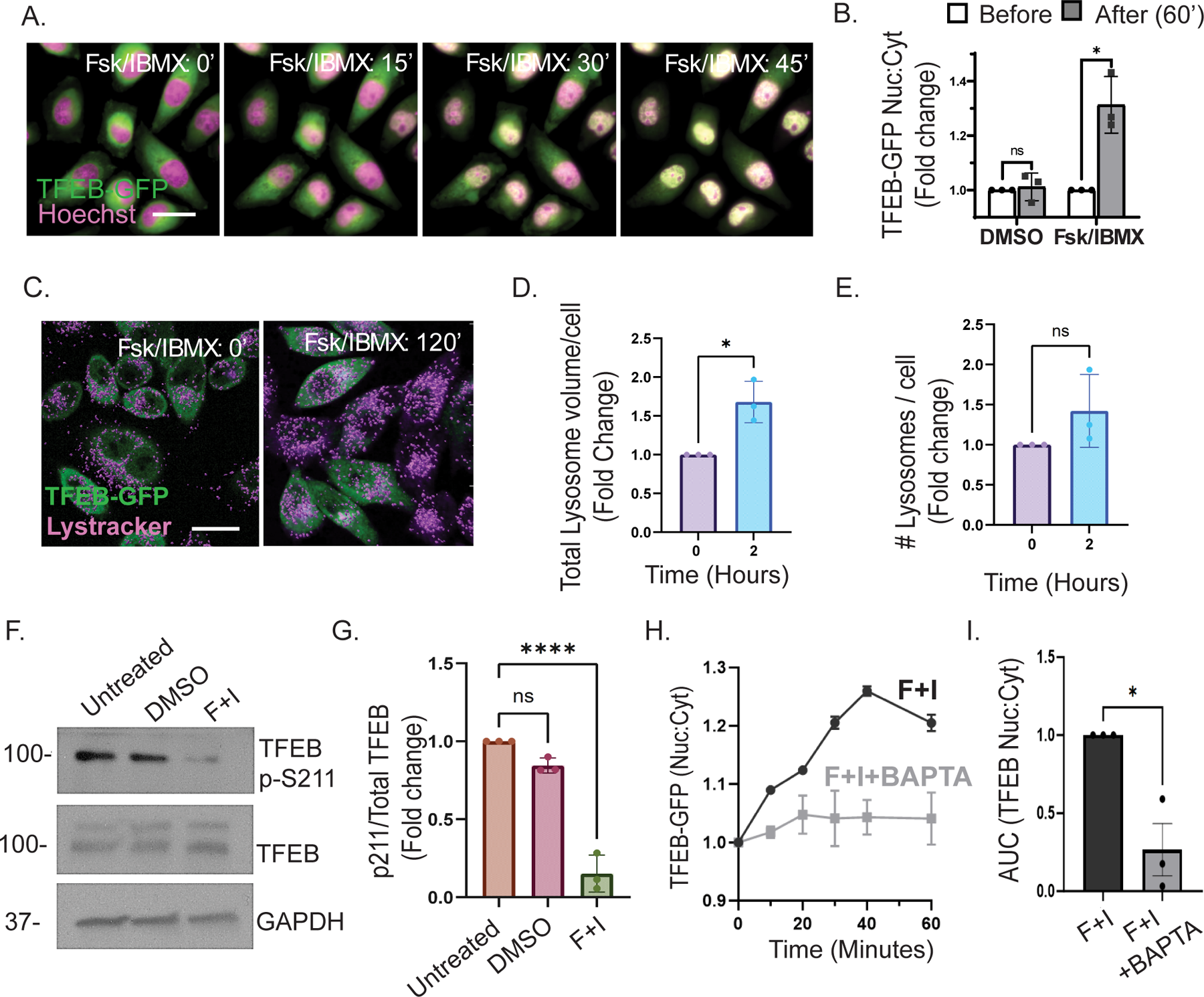
cAMP stimulates TFEB nuclear import via calcium-dependent signaling and enhances lysosomal volume. (A) Fluorescence microscopy of HeLa cells stably expressing TFEB-GFP treated with cAMP agonists (10 µM Fsk + 0.5 mM IBMX) shows the rapid redistribution of TFEB-GFP from the cytoplasm to the nucleus. (B) Quantification of the nuclear-to-cytoplasmic TFEB-GFP ratio resulted in a significant increase in nuclear import within 60 minutes, compared to the DMSO control (n=3, paired T-test *p < 0.05). (C) Confocal imaging reveals enhanced lysosomal volume in HeLa cells, as evidenced by increased LysoTracker Deep Red staining following cAMP treatment. (D-E) Quantification of confocal micrographs in panel (C) shows that cAMP treatment for 2 hours significantly increases total lysosome volume per cell, while the number of lysosomes per cell (E) remains unchanged compared to control (n=3, ratio paired t test, two-tailed test *p < 0.05). (F) Western blot analysis of whole-cell lysates from HeLa cells treated with DMSO (30 min), Fsk/IBMX (30 min), or left untreated. Blots were probed with a phospho-specific antibody against TFEB S211. GAPDH served as a loading control. (G) Densitometric quantification of phospho-S211 signal using ImageJ, normalized to GAPDH, and analyzed in GraphPad Prism. A significant decrease in S211 phosphorylation was detected upon Fsk/IBMX treatment compared to DMSO control (n=3, one-way ANOVA, Tukey’s multiple comparison test ****p < 0.01). (H) Quantification of the nuclear-to-cytoplasmic TFEB-GFP ratio at 0, 20, 40, and 60 minutes following Fsk/IBMX treatment in calcium-deficient medium, with or without the calcium chelator BAPTA-AM. Calcium chelation markedly reduces the nuclear localization of TFEB over time. (I) Area under the curve (AUC) analysis of the nuclear-to-cytoplasmic ratio panel (H) further demonstrates a significant decrease in nuclear TFEB accumulation in the presence of BAPTA-AM (n = 3, paired t-test *p < 0.05).

TFEB nuclear import is known to be triggered by Ca²⁺-dependent dephosphorylation of S211(Tong and Song, 2015). Since cAMP can increase intracellular Ca²⁺ (Akerman et al., 2024), we examined TFEB S211 phosphorylation by western blot analysis, which revealed an ∼80% reduction in phospho-S211 after 30 minutes of Fsk/IBMX treatment (Fig. 1F-G). To test whether this response depends on Ca²⁺, Fsk/IBMX treated cells were imaged in Ca²⁺-free medium with or without the intracellular Ca²⁺ chelator BAPTA-AM (10 μM). TFEB-GFP nuclear import was significantly inhibited in Ca²⁺-chelated conditions, as shown by reduced nuclear-to-cytoplasmic ratios over time and reduced area-under-the-curve measurements (Fig. 1H-I). AML-12 cells displayed a similar calcium requirement (Fig. S1C). Interestingly, dephosphorylation of S211 still occurred in the presence of BAPTA-AM (Fig. S1D-E), indicating that cAMP suppresses S211 phosphorylation independently of intracellular Ca²⁺, while Ca²⁺ is required for nuclear import itself. We also assessed whether cAMP affects phosphorylation at S122, another mTOR-dependent regulatory site, and found no detectable changes (Fig. S1F-G), in contrast to the robust decrease at S211. Together, these results suggest that cAMP promotes TFEB nuclear import through calcium signaling and the selective dephosphorylation at S211, leading to increased lysosomal volume.

### PKA inhibition prevents TFEB nuclear export

Next, we sought to determine the impact of PKA activity on cAMP-mediated TFEB nuclear import/export dynamics. To do this, we performed western blot analysis of nuclear and cytoplasmic fractions as well as high-throughput fluorescence microscopy of HeLa cells stably expressing TFEB-GFP in parallel. As shown in the representative western blot (Fig. 2A), TFEB underwent a rapid but transient nuclear accumulation following cAMP stimulation, peaking between 0.5 and 1 hours before gradually redistributing to the cytoplasm between 2 and 8 hours. Strikingly, pre-treatment with the PKA inhibitor H89 had no effect on acute TFEB import but appeared to block subsequent export resulting in sustained nuclear TFEB levels throughout the 8-hour time course (Fig. 2B). Note that, in our hands, TFEB was detected as a doublet by western blot, but both bands correspond to TFEB by mass spec analysis, and their differential migration is perhaps consistent with phosphorylation-dependent mobility changes by SDS-PAGE. Next, we used high-throughput fluorescence microscopy to quantitatively assess TFEB subcellular localization dynamics in a large number of cells. Using CellProfiler, “nucleus” was defined using Hoechst staining while “cytosol” was defined as a ∼8.5 µm perinuclear band surrounding each nuclear mask. TFEB-GFP intensity was measured within both regions to generate nuclear-to-cytoplasmic ratios at each time point. Approximately 15,000 cells were analyzed per condition (n=3 independent experiments). As shown in representative images and quantification from (Fig. 2C-D), this microscopy-based quantification supported the western blot findings and confirmed a ∼50% increase in nuclear: cytoplasmic TFEB-GFP at 0.5 hours post-Fsk/IBMX treatment, followed by progressive nuclear export at 2 and 8 hours. Notably, H89 treatment prevented this export, resulting in a sustained 2-fold increase in nuclear TFEB-GFP throughout the time course. Together, these results suggest that PKA activity is not required for TFEB nuclear import but is essential for TFEB export.

**Figure 2:**
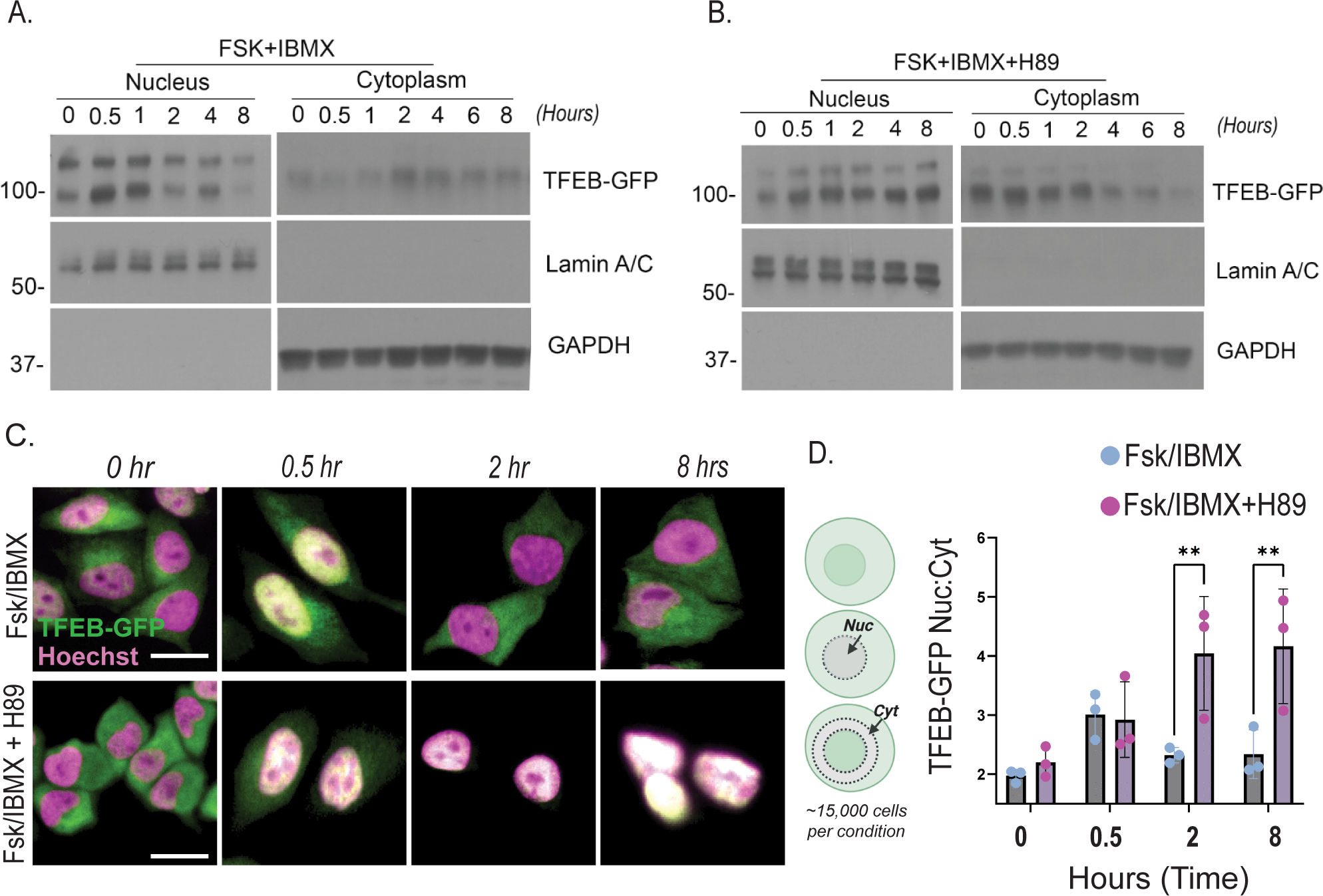
cAMP promotes transient TFEB nuclear import, and PKA inhibition prevents TFEB nuclear export. (A) HeLa cells expressing TFEB-GFP were treated with cAMP agonists (10 µM Fsk + 0.5 mM IBMX) with or without the PKA inhibitor H89 (50 µM). Western blot analysis shows a transient nuclear import of TFEB between 0–1 hour, followed by nuclear export from 2–8 hours after Fsk/IBMX treatment. (B) In the presence of H89, nuclear export is inhibited, and TFEB remains nuclear from 2–8 hours post-treatment. (C–D) In parallel with the western blot (n = 2), fluorescence microscopy was used to visualize TFEB dynamics at the single-cell level. To quantitatively assess TFEB dynamics across a larger population, high-throughput imaging was performed in three independent experiments (∼15,000 cells per condition). Nuclei were segmented using Hoechst staining, and cytosolic regions were defined as an 8.5 µm perinuclear band using CellProfiler. TFEB–GFP fluorescence intensity was measured in both compartments to calculate nuclear-to-cytoplasmic (N/C) ratios at 0, 0.5, 2 and 8 hours. Representative micrographs (C) and quantification of the nuclear-to-cytoplasmic TFEB-GFP ratio (D) confirm nuclear import at 0.5 hours and export between 2–8 hours after Fsk/IBMX treatment. H89 treatment prevents this export, resulting in sustained nuclear TFEB localization. Statistical analysis was performed using two-way ANOVA with Sidak’s post hoc test (**p < 0.01).

### PKA inhibition prevents TFEB nuclear export independent of Fsk/IBMX treatment

TFEB is a key stress-responsive transcription factor that becomes activated and translocates to the nucleus under nutrient starvation. To further assess the role of PKA activity in TFEB nuclear export, we employed a TFEB export assay as previously described(Li et al., 2018; Napolitano et al., 2018; Ruolo et al., 2025), whereby cells are first subjected to 1.5 hours nutrient starvation using HBSS to inhibit mTOR and trigger TFEB nuclear translocation, followed by nutrient refeeding for 8 hours to stimulate TFEB nuclear export back to the cytoplasm. Refeeding was done in the presence of H89 (50 μM) or vehicle control (DMSO). As shown in fluorescence micrographs in (Fig. 3A), DMSO control cells exhibit normal TFEB-GFP nuclear entry upon starvation, as well as the expected nuclear export upon nutrient re-feeding. In contrast, cells treated with H89 still underwent starvation-induced nuclear import, however TFEB was retained in the nucleus upon refeeding, indicating impaired nuclear export despite nutrient availability (Fig. 3B). Quantitative analysis of nuclear TFEB fluorescence intensity confirmed significantly higher nuclear retention in H89-treated cells compared to controls (Fig. 3C). These findings suggest that PKA activity plays a critical role in promoting TFEB nuclear export upon nutrient refeeding, independent of direct cAMP stimulation.

**Figure 3:**
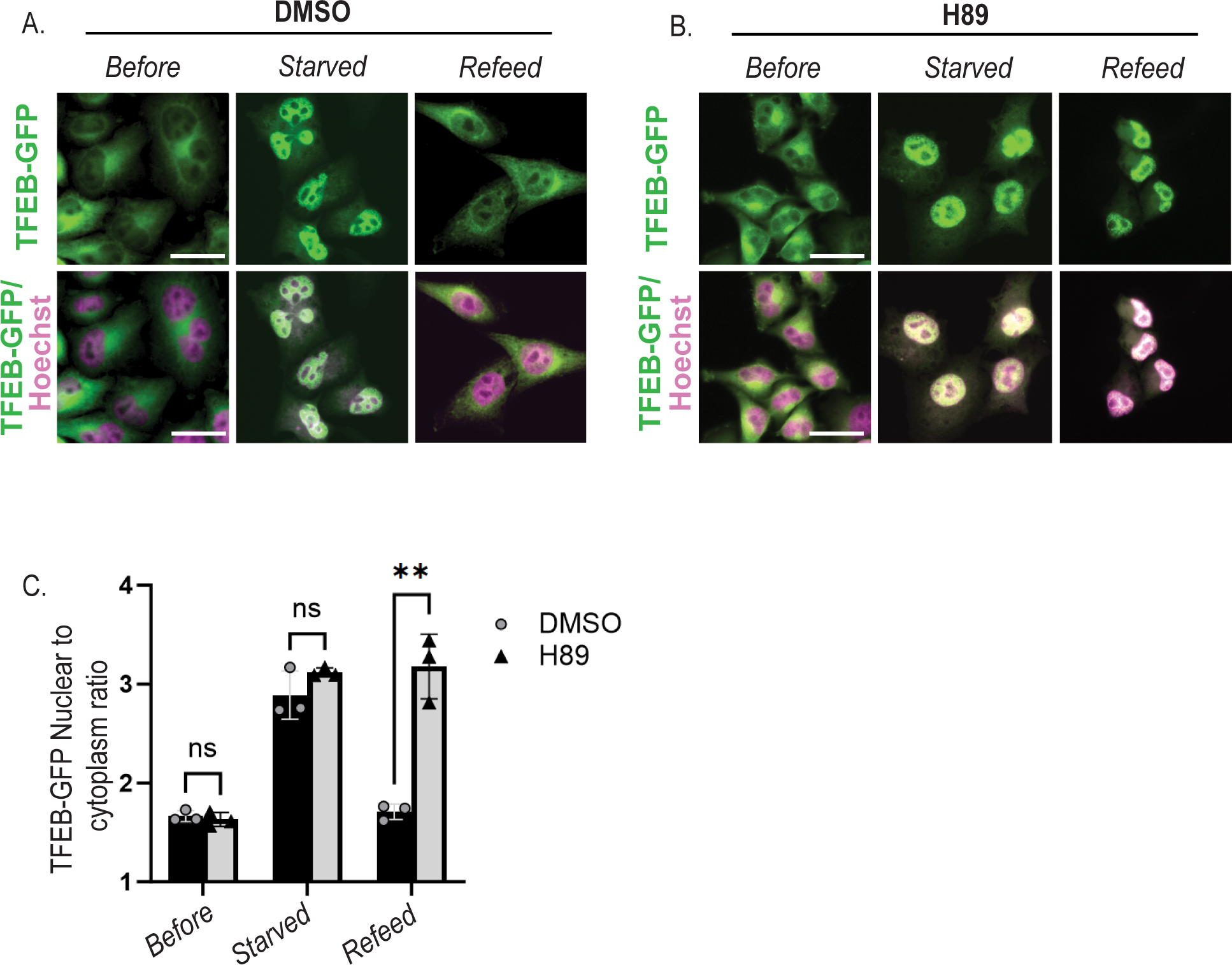
PKA inhibition prevents TFEB nuclear export independent of Fsk/IBMX treatment. (A) Fluorescence micrographs of DMSO-treated HeLa cells expressing TFEB–GFP show that TFEB translocates to the nucleus within 30 min of HBSS-induced starvation but relocates to the cytoplasm 8 h after refeeding, indicating nuclear export. (B) In contrast, treatment with the PKA inhibitor H89 (50 µM) prevents TFEB export, resulting in sustained nuclear localization after 8 h of refeeding under identical conditions. (C) Quantification of TFEB–GFP fluorescence confirms significantly greater nuclear retention in H89-treated cells compared with DMSO controls following starvation-induced nuclear import and subsequent refeeding (n = 3, t-test **p < 0.01).

### cAMP and PKA differentially regulate TFEB target gene transcription

To assess TFEB transcriptional activity in response to cAMP signaling, we measured the expression of a panel of TFEB target genes that contain CLEAR motifs in their promoter regions. HeLa cells were treated with Fsk+IBMX alone or in combination with H89 for 2 and 8 hours, with untreated cells serving as controls. Total RNA was extracted, and mRNA levels of TFEB target genes(Palmieri et al., 2011)—LAMP1, MCOLN1, CTSD, CLN7, and ATP6V0E1—were quantified using qPCR. Gene expression was normalized to GAPDH, and relative fold changes were calculated using the ΔΔCt method. At the 2-hour time point, Fsk/IBMX treatment induced a significant upregulation of TFEB target genes, indicating activation of TFEB-mediated transcription in response to acute cAMP stimulation. However, by 8 hours, expression of these target genes declined to near-baseline levels, suggesting that the cAMP-induced transcriptional response is transient (Fig. 4A). In contrast, pretreatment with the PKA inhibitor H89 resulted in sustained elevation of TFEB target gene expression at both 2 and 8 hours of Fsk/IBMX treatment (Fig. 4B). These findings suggest that PKA activity acts to constrain the duration of TFEB-driven transcription during prolonged cAMP signaling, potentially by promoting TFEB nuclear export at later time points.

**Figure 4:**
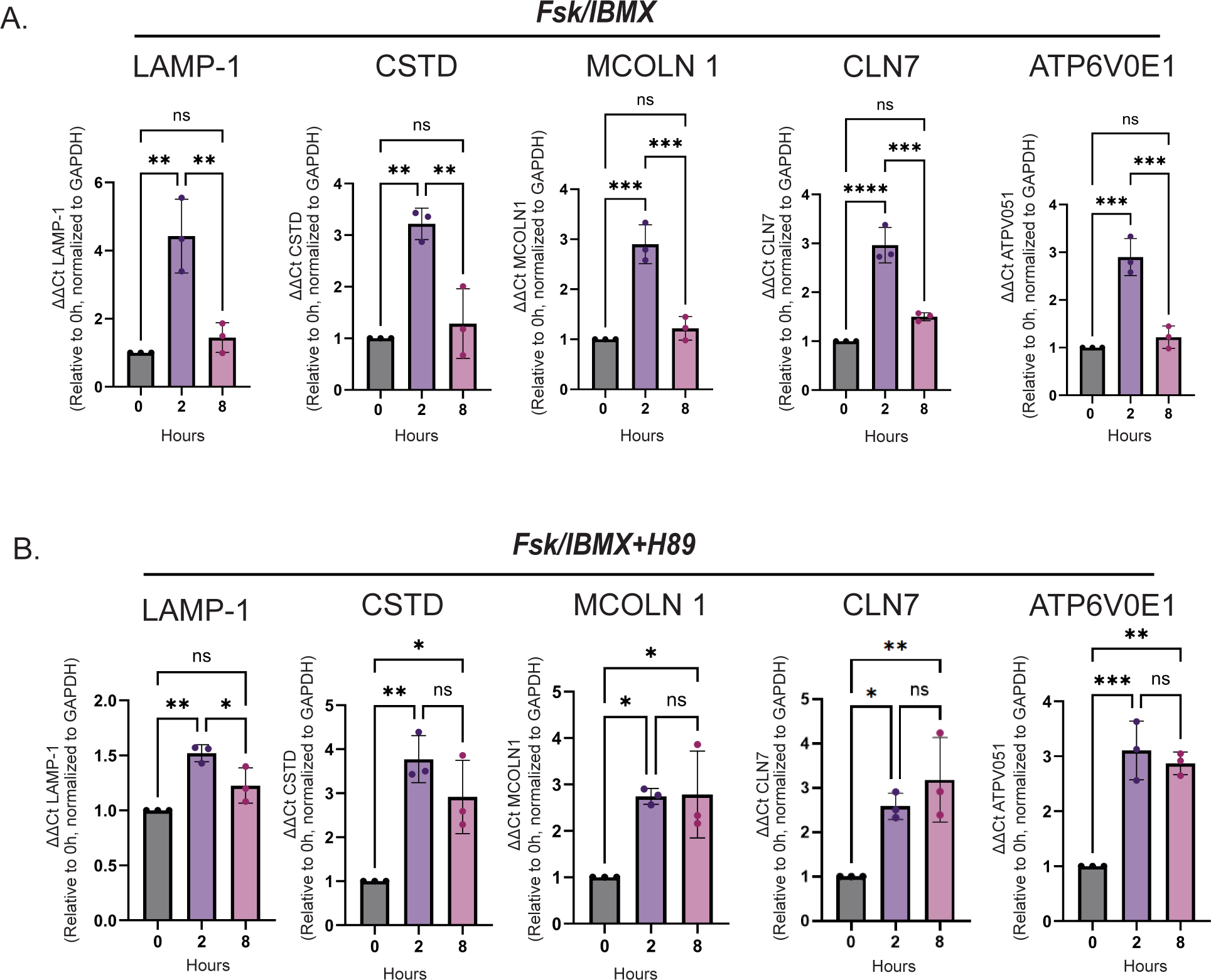
cAMP and PKA differentially regulate TFEB target gene transcription. HeLa cells were treated with Fsk/IBMX or combined with the PKA inhibitor H89 for 2 and 8 hours. Total RNA was extracted, and mRNA levels of TFEB target genes (LAMP1, MCOLN1, CTSD, CLN7, *ATP6V0E1*) were quantified by qPCR. Gene expression was normalized to the housekeeping gene GAPDH, and relative fold changes were calculated using the ΔΔCt method. (A) Fsk/IBMX treatment induced a significant upregulation of TFEB target gene expression at 2 hours; however, by 8 hours, expression levels declined markedly, indicating a transient transcriptional response to cAMP signaling. (B) In contrast, co-treatment with the PKA inhibitor H89 led to a significant increase in TFEB target gene expression at both 2 and 8 hours (n=3, one-way ANOVA, Tukey’s multiple comparison test, *p < 0.05, **p < 0.01, ***p < 0.001), suggesting that PKA activity tunes the duration of TFEB-mediated transcription during prolonged cAMP stimulation.

### Differential phosphorylation of TFEB during nuclear import vs export phases

To investigate the effect of cAMP stimulation on the overall phosphorylation profile of TFEB, we conducted phosphoproteomic analysis of TFEB-GFP in HeLa cells treated with Fsk/IBMX (noted F+I), with or without PKA inhibition by H89 for 8 hours (export phase). GFP-Trap was used to purify TFEB-GFP, followed by mass spectrometry. As shown in the phospho-peptide heatmap in Figure 5A, we identified a significant increase in phosphorylation of S467 and the adjacent residue S466 in F+I treated cells, but this increase was not observed in H89-treated or F+I+H89-treated cells. As these residues fall within PKA phosphorylation consensus sequence (RRxS), our results suggest that S466 and S467 may facilitate PKA-dependent TFEB nuclear export, consistent with our findings that PKA inhibition prevents export upon cAMP-mediated import (Fig. 2,3). We next examined TFEB localization in cells transiently expressing WT TFEB-GFP, a phospho-null mutant (S466,467A), or a phospho-mimetic mutant (S466,467D). WT TFEB underwent nuclear import at 30 min and cytoplasmic relocalization by 8 hours of Fsk/IBMX treatment. The phospho-mimetic mutant behaved similarly. In contrast, TFEB(S466,467A) was retained within the nucleus at both 30 min and 8 h (Fig. 5B,C), underscoring the critical role of these residues in TFEB nuclear export. To evaluate individual contributions of S466 and S467, we analyzed single-site phospho-null mutants S466A and S467A, as well as single-site phospho-mimetic mutants S466D and S467D. All single mutants underwent similar nuclear import (30 mins) and export to cytoplasm (8h) upon Fsk/IBMX treatment similar to wild-type TFEB (Fig. S2A–B). These results indicate that phosphorylation of both S466 and S467 cooperatively promote nuclear export downstream of cAMP/PKA signaling. We also performed phosphoproteomic analysis during the import phase (30 min) identified increased cAMP-induced phosphorylation of S109, S114, and S122. However, neither phospho-null nor phospho-mimetic mutations at these sites altered TFEB localization at baseline or after cAMP stimulation (Fig. S3A–F), consistent with previous work showing that these residues contribute to cytoplasmic retention primarily in conjunction with S211(Martina and Puertollano, 2018). Together, these data identify S466 and S467 as key PKA-regulated sites required for TFEB nuclear export.

**Figure 5:**
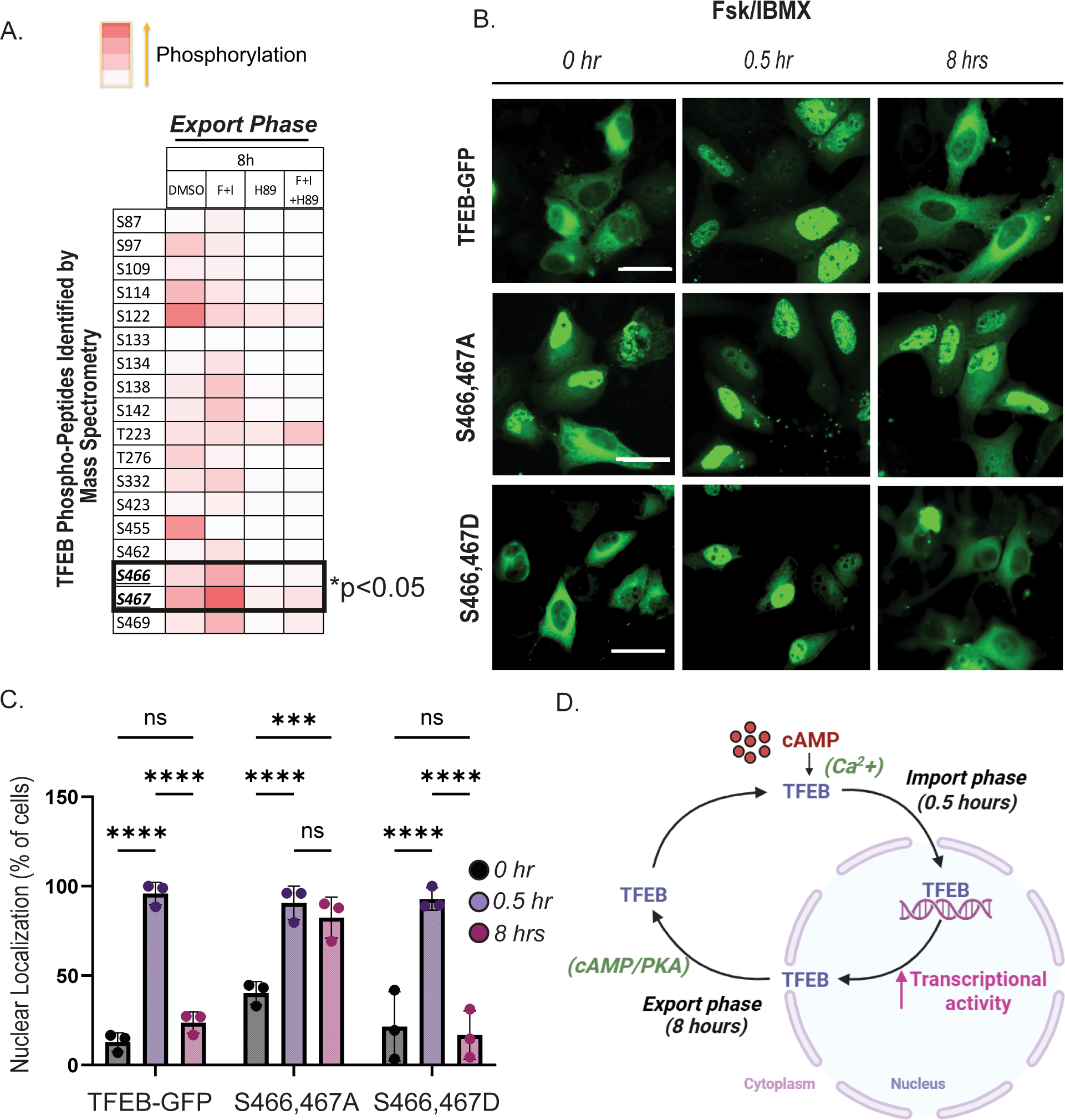
Differential phosphorylation of TFEB during nuclear import vs export phases: (A) Heatmap of TFEB–GFP phosphopeptides identified by GFP-Trap pulldown from HeLa cells treated for 8 h with DMSO, Fsk (10 µM) + IBMX (0.5 mM), Fsk/IBMX + H89, or H89 alone (50 µM), analyzed by mass spectrometry (n = 3 independent experiments). Phosphorylation at S466 and S467 were significantly increased during the nuclear export phase in a PKA-dependent manner. (B) Fluorescence microscopy of HeLa cells expressing wild-type TFEB or phospho-null (S466,467A) and phospho-mimetic (S466,467D) mutants. (C) Quantification of nuclear TFEB–GFP levels (n = 3; 100 cells per condition) shows enhanced nuclear retention of the phospho-null mutant and reduced nuclear localization of the phospho-mimetic mutant, comparable to wild-type TFEB(n = 3; 100 cells/condition; Two-way ANOVA, Sidak’s multiple comparison test; *p* < 0.05), supporting a critical role of phosphorylated serine residues 466 and 467 in regulating TFEB nuclear export. (D) Model depicting cAMP/PKA-dependent regulation of TFEB nucleocytoplasmic shuttling, in which cAMP promotes TFEB nuclear import, whereas PKA activity drives its nuclear export to tune its transcriptional activity.

## Discussion

While TFEB nuclear import has been extensively studied, mechanisms governing its nuclear export have only recently begun to receive focused attention. Our study uncovers a novel mechanism by which cAMP regulates the subcellular localization and transcriptional activity of the master lysosomal regulator TFEB through calcium- and PKA-dependent signaling events. We reveal a previously unrecognized phase-specific regulation of TFEB: a rapid, calcium-dependent nuclear import (Fig. 1) independent of PKA activity, followed by a delayed, PKA-dependent nuclear export (Fig. 2,3) that appears to be regulated by phosphorylation at serine residues S466 and S467 (Fig. 5). These findings highlight a new role for cAMP-PKA signaling in tuning transcriptional programs essential for lysosomal homeostasis (Fig. 4).

### Tuning lysosome dynamics through transient TFEB activation

TFEB functions as a dynamic metabolic regulator, shuttling between the nucleus and cytoplasm in response to extracellular stimuli. We show that cAMP-induced TFEB nuclear import is rapid (30–60 min) but transient, with efficient nuclear export occurring between 2 and 8 hours. This suggests a mechanism to tune the duration of TFEB transcriptional activity. Such transient activation may be important during nutrient refeeding, when lysosomal function must return to baseline after starvation, or during the cell cycle, where excess lysosomal biogenesis could interfere with mitotic progression(Contreras and Puertollano, 2023; Franco-Juarez et al., 2022). Fine-tuning lysosome biogenesis enables reversible adaptation without the energetic or structural cost of sustained activation.(Mutvei et al., 2023; Yang and Wang, 2021; Zoncu and Perera, 2022). PKA is a known transcriptional regulator, most notably by facilitating CREB-mediated transcription(Zhang et al., 2020) and also as a modulator of mTORC1 signaling(Jewell et al., 2019). Prolonged TFEB activation can be deleterious; constitutive TFEB activity has been linked to tumorigenesis(Follo et al., 2019; Gwak et al., 2022; Yun et al., 2021). The role of PKA in cancer is context-dependent: elevated PKA can promote oncogenic programs (Na et al., 2020; Chan et al., 2023; Shirani et al., 2024), whereas suppression of PKA signaling, including through overexpression of protein kinase inhibitor beta (PKIB), a competitive inhibitor that binds the PKA catalytic subunit, can also enhance tumor progression (Liu et al., 2025). Because PKA signaling is spatially organized by AKAP scaffolds (Rosenthal et al., 2024), both hyperactivation and inhibition can contribute to oncogenesis depending on subcellular context (Chan et al., 2023; Liu et al., 2025; Na et al., 2020; Rosenthal et al., 2024; Shirani et al., 2024). In this framework, PKA-dependent TFEB nuclear export provides a mechanism to activate lysosomal gene expression transiently while preventing prolonged TFEB activity. This tiered regulation may represent a general strategy by which PKA shapes transcriptional responses—activating them briefly to meet immediate demands and ensuring timely termination to maintain cellular homeostasis.

### cAMP-Induced TFEB nuclear import is calcium-dependent but PKA-independent

Our data show that cAMP-induced TFEB nuclear import is independent of PKA but critically dependent on calcium. Phosphoproteomic analysis revealed increased phosphorylation of S109, S114, and S122 in a PKA-dependent manner (Fig. 5A), which do not contribute to cAMP-mediated import (Fig. S3B-C). While the residue S211, a key mTOR regulatory site, was not detected by mass spectrometry, western blotting showed reduced S211 phosphorylation upon cAMP stimulation (Fig. 1F-G). Interestingly, Ca²⁺ chelation with BAPTA-AM blocked nuclear import despite continued S211 dephosphorylation (Fig. S1H-I), suggesting additional calcium-dependent mechanisms facilitate TFEB nuclear entry. The source of this calcium may involve multiple known cAMP–Ca²⁺ coupling mechanisms. Firstly, cAMP can elevate intracellular calcium levels is through modulation of the inositol 1,4,5-trisphosphate (IP₃) receptor (IP₃R), a calcium channel located on the membrane of the endoplasmic reticulum (ER)(Taylor, 2017). This cAMP-IP₃R-Ca²⁺ axis is especially relevant in cells with highly compartmentalized signaling, such as neurons(Zhang et al., 2012). Secondly, cAMP can also signal through EPAC, independently of PKA(Lezcano et al., 2018; Ruiz-Hurtado et al., 2013), to regulate calcium levels by affecting channels like ryanodine receptors (RyRs) and store-operated calcium entry (SOCE)(Lezcano et al., 2018; Yip et al., 2023). Additionally, in some contexts, cAMP–EPAC signaling can activate phospholipase C epsilon (PLC-ε), leading to increased IP₃ production and subsequent IP₃R-mediated calcium release from the ER (Oestreich et al., 2009; Oestreich et al., 2007). Together, these pathways provide multiple routes for cAMP to influence intracellular calcium signaling, supporting the regulation of dynamic TFEB.

### cAMP/PKA dependent phosphorylation controls TFEB nuclear export

Our study identifies cAMP/PKA signaling as a key regulator of TFEB nuclear export. We demonstrate that phosphorylation at S466 and S467 is required for nuclear export, as alanine substitution at these residues sustains TFEB in the nucleus following cAMP stimulation. These sites lie within an RRSS motif, consistent with direct PKA phosphorylation, though we cannot exclude the possibility that PKA could modulate TFEB through another intermediate kinase. Regardless, inhibition of PKA abolishes nuclear export, establishing a PKA-dependent mechanism. The additive effect of the S466A/S467A double mutant over the single S466A or S467A mutants further underscores the cooperative role of these adjacent serine residues.

Nuclear retention of S466,S467A TFEB (Fig. 5B-C) could suggest a nuclear PKA pool, potentially anchored by nuclear A-kinase anchoring proteins such as AKAP8 or AKAP8L(Clister et al., 2019; Wang et al., 2025). An interesting future direction will be to probe the role of nuclear AKAP–PKA microdomain as a plausible spatial platform for TFEB regulation. S466 and S467 are also important for TFEB transcriptional activity, as AMPK phosphorylation at S466, S467 and S469 enhances TFEB-DNA interaction independently of nuclear localization(Paquette et al., 2021). Thus, phosphorylation at S466 and S467 may act as a temporal switch to tune lysosomal and metabolic programs.

Although the current study establishes a possible framework linking cAMP/PKA activity to TFEB dynamics, several questions remain. Future studies will be required to further define the cAMP–calcium import mechanism, explore additional kinases contributing to S466 and S467 phosphorylation, and investigate the spatial relationship between TFEB and nuclear AKAP–PKA complexes to reveal potential nuclear export pathways and strategies to modulate TFEB activity via nuclear signaling microdomains.

## Methods and Materials

### Cell culture and transfection

Parental HeLa cells were acquired from ATCC (Cat. No. CCL-2), and HeLa cells stably expressing TFEB-GFP were a kind gift from Richard Youle (NIH)(Nezich et al., 2015). Cells were cultured in DMEM containing 4.5 g/L glucose (Thermo Fisher, Cat. No. 10-013-CV) or Ca²⁺-free DMEM (Gibco, Cat. No. 21068028), supplemented with 10% fetal bovine serum (Gibco, Cat. No. 26140079) and 1% penicillin-streptomycin (Gibco, Cat. No. 15140122). Cells were maintained at 37 °C in a humidified incubator with 5% CO₂. Pharmacological treatments included Forskolin (10 µM; Sigma-Aldrich, Cat. No. F6886) and IBMX (0.5 mM; Sigma-Aldrich, Cat. No. I7018), were added directly to the culture medium for periods ranging from 30 minutes to 8 hours. Cells were pre-treated for 30 minutes with the PKA inhibitor H89 (50 µM; Sigma, Cat. No. B1427) or the Ca²⁺ chelator BAPTA-AM (10 µM; Cayman Chemical, Cat. No. 15551-5) before stimulation with Forskolin/IBMX.

TFEB-GFP phosphorylation site mutants (S467A, S467E, and S466,S467A) were generated by site-directed mutagenesis using a commercial kit (NEB, Cat. No. E0554S) and confirmed by sequence analysis. TFEB-GFP S109,114,122A and S109,114,122E were commercially generated by Vectorbuilder. For all constructs, HeLa cells were transfected at 60–80% confluency using Lipofectamine™ 3000 (Invitrogen, Cat. No. L3000015) following the manufacturer’s instructions.

### Live-cell microscopy

HeLa cells stably expressing TFEB-GFP were seeded into 35mm glass bottom imaging dishes (Cat. No. Cell E&G, GBD00004-200) and maintained in FluoroBrite™ DMEM (Thermo Fisher Scientific, Cat. No. A1896701). Lysosomes were visualised using LysoTracker™ Deep Red (Invitrogen, Cat. No. L12492) at a 1:20,000 dilution for 30 minutes at 37 °C. Live-cell confocal imaging was performed using a BC43 Benchtop Confocal Microscope (Andor, Oxford Instruments) equipped with a 40X oil immersion objective and an environmental chamber maintained at 37 °C with 5% CO₂. TFEB-GFP and LysoTracker fluorescence were excited with 488 nm and 640 nm lasers, respectively. Lysosome volume and number were quantified using Imaris software (version 10.2.0) based on 3D surface rendering and volumetric thresholding.

To monitor TFEB nuclear import, time-lapse live-cell imaging was conducted using a Nikon Eclipse Ti2 inverted fluorescence microscope with a 20X air objective. HeLa cells stably expressing TFEB-GFP were imaged every 5 minutes for 60 minutes following treatment with forskolin and IBMX. For GFP emission, a 488 nm LED was used to capture the subcellular localization of TFEB. Cells were maintained under physiological conditions (37 °C, 5% CO₂) throughout the imaging time course.

### High-content fluorescence microscopy

HeLa cells were seeded on sterile glass coverslips or in glass-bottom 96-well μ-plates (IBIDI, Cat. No. 89626) at a density of 2,000 cells per well. Following treatment, cells were washed with phosphate-buffered saline (PBS; ThermoScientific, Cat. No. MT21040CV) and fixed in 3% formaldehyde for 20 minutes at room temperature. Nuclei were stained with Hoechst 33342 (Invitrogen, Cat. No. H3570) for 30 minutes, followed by PBS washes.

Fluorescence imaging was performed using a Nikon Eclipse Ti2 inverted microscope equipped with a 20X air objective and DAPI and GFP filter sets. DAPI and GFP fluorescence were excited using 359 nm and 488 nm LEDs respectively. Nuclear masks were generated from Hoechst staining using CellProfiler (version 4.2.8), and cytoplasmic masks were defined as rings surrounding each nucleus. The cytoplasmic ring was defined by expanding the nuclear mask outward by 16 pixels and then shrinking it by 3 pixels, resulting in a 13-pixel-wide band (∼8.5 µm based on image resolution).TFEB-GFP fluorescence intensity was quantified in both nuclear and cytoplasmic regions in the GFP channel. Subcellular localization of TFEB was assessed by calculating the nuclear-to-cytoplasmic intensity ratio per cell. For each condition and time point, seven non-overlapping high-resolution fields were captured per well. Quantification was based on at least 15,000 cells per condition across three independent biological replicates.

### Western blotting

Subcellular fractionation was performed using the REAP (Rapid, Efficient, and Practical) method as previously described(Suzuki et al., 2010). In brief, cells were lysed in ice-cold 0.1% NP-40 in PBS to release the cytoplasmic contents, followed by brief centrifugation to separate the cytoplasmic fraction. The remaining pellet was washed once with 0.1% NP-40 to remove residual cytoplasmic proteins and then resuspended to obtain the nuclear fraction. This method allows rapid and gentle separation of cytoplasmic and nuclear proteins for subsequent analysis(Suzuki et al., 2010). Whole cell lysates were collected using 1X RIPA buffer, supplemented with Protease inhibitor cocktail tablets cOmplete, EDTA-free (Sigma, Cat. No. 11836170001) and Halt phosphatase inhibitor (VWR, Cat. No. PI78427). Lysates were sonicated and centrifuged at 16,000 × *g* for 10 minutes at 4°C to pellet cellular debris.

Protein concentration was measured using the Pierce BCA Protein Assay Kit (Thermo Fisher Scientific, Cat. No. 23227), following the manufacturer’s instructions. Equal amounts of protein (20–40 µg) were resolved on 4-20% SDS-PAGE gels and transferred to PVDF membranes (Millipore, Cat. No. IPVH00010) using a semi-dry transfer system. Membranes were incubated overnight at 4°C with the following primary antibodies: TFEB (1:1000, CST, Cat. No. 37785S), p-S211 TFEB (1:1000, CST, Cat. No. E9S8N), GFP (1:1000, Chromotek, Cat. No. PABG1-100), Lamin A/C (1:1000, CST, Cat. No. 4777S) – nuclear loading control, GAPDH (1:5000, CST, Cat. No. 5174S) – cytoplasmic/whole-cell loading control. After three washes with TBST, membranes were incubated with HRP-conjugated secondary antibodies (1:25,000, Invitrogen), Rabbit (Cat. No. G21234) and Mouse (Cat. No. G21040) for 30 minutes at room temperature. Bands were visualized using ECL detection reagent (ThermoFisher Scientific, Cat. No. 34580) and developed using a film-based imaging system. Densitometric analysis of western blots was performed using ImageJ.

### Quantitative Real-Time PCR (qPCR)

Total RNA was extracted from HeLa cells using the RNeasy Mini Kit (Qiagen, Cat. No. 74104) according to the manufacturer’s protocol, including an on-column DNase I digestion step (Qiagen, Cat. No. 79254) to eliminate residual genomic DNA. RNA concentration and purity were assessed using a NanoDrop one C spectrophotometer (Thermo Fisher Scientific), and RNA integrity was verified by electrophoresis or standard 260/280 and 260/230 ratios.

For one-step RT-qPCR, 1 µg of total RNA was used as the template in a 20 µL reaction using the Luna® Universal One-Step RT-qPCR Kit (New England Biolabs, Cat. No. E3005L), following the manufacturer’s instructions. Reactions were performed on a Bio-Rad CFX96 Touch™ Real-Time PCR Detection System, and amplification was monitored using the SYBR® Green detection chemistry provided in the kit. Target genes included LAMP1, CLN7, MCOLN1, ATP6V0E1, and CTSD, with GAPDH as the normalization control. Primer sequences were derived from previously published work (Li et al., 2019; Zeng et al., 2017) as follows(Li et al., 2019b; Zeng et al., 2017): Primer sequences were: LAMP1 forward, TCTCACTGAACTACCACACCA; reverse, AGTGTATGTCCTCTTCCAAAAGC.

CTSD forward, CTGTGGAGGACCTGATTGC; reverse, CTGGACTTGTCGCTGTTGTAC. MCOLN1 forward, GCCGGTATGCGTATGTCC; reverse, TTTGAGCGTGAGGTTCTTGTA. CLN7 forward, CTGTTCACGCTGGTCTACTTCT; reverse, TCCCATCAGGGCGTATTT. ATP6V0E1 forward, GCCTTGGTTCATCCCTAA; reverse, AGGCCAATGATACTTCAGATA. The thermal cycling conditions were as follows: one denaturation cycle at 95°C for 2 minutes, forty cycles of 95°C for 15 seconds, one cycle at 60°C for 30 seconds, and melt curve analysis from 65°C to 95°C to confirm amplicon specificity. Relative gene expression levels were calculated using the 2^–ΔΔCt method. All reactions were performed in technical triplicate, and results represent at least three independent biological replicates.

### GFP-Trap immunoprecipitation and phospho-proteomic analysis

HeLa cells stably expressing TFEB-GFP were lysed, and TFEB-GFP was immunoprecipitated using GFP-Trap agarose beads (Chromotek, Cat. No. gta-20) following the manufacturer’s protocol. Eluted proteins were separated by SDS-PAGE using Criterion XT Pre-cast gels (Bio-Rad, Cat. No. 3450117) and XT MOPS running buffer (Bio-Rad, Cat. No. 1610788). Protein bands were visualized by Coomassie Brilliant Blue G-250 staining, and bands of interest were excised for downstream analysis. In-gel digestion was performed using the Pierce In-Gel Tryptic Digestion Kit (Thermo Scientific, Cat. No. 89871). Gel slices were reduced with dithiothreitol, alkylated with iodoacetamide, digested overnight with trypsin at 37°C, and peptides were extracted and reconstituted in 5% acetonitrile and 0.05% trifluoroacetic acid. Samples were analyzed by nanoLC-MS/MS using a 2-hour gradient on a 0.075 mm × 250 mm C18 column coupled to an Orbitrap Eclipse mass spectrometer (Thermo Scientific) operating in OT-OT-HCD mode. MS/MS data were searched using Mascot (v2.7.0, Matrix Science) against the UniProt human reference proteome (UP000005640, release 2023-11-20) and a common contaminant database (cRAP_20150130.fasta). Parameters included: trypsin digestion, 15 ppm precursor tolerance, 0.060 Da fragment ion tolerance, and variable modifications for carbamidomethyl (C), oxidation (M), deamidation (N/Q), and phosphorylation (S/T/Y). Peptide and protein identification were validated in Scaffold (v5.3.2, Proteome Software). Peptide identifications were accepted at >80% probability using Peptide Prophet algorithm (Keller, A et al Anal. Chem. 2002;74(20):5383-92) with delta-mass correction, and protein identifications at >99% probability with at least two unique peptides (Protein Prophet).

### Quantitative proteomics and phosphosite analysis

Protein quantification was performed using Proteome Discoverer (Thermo Fisher Scientific, v2.4). MS/MS data were searched using the same parameters and databases as described above. Peptide validation was performed using Percolator with a posterior error probability (PEP) threshold of 0.01. A decoy database search strategy was used to ensure a 1% false discovery rate (FDR) for high-confidence identifications.

Quantification was based on precursor ion intensity, and peptide abundances were normalized to total peptide amount. Scaled abundances were adjusted so that the average for each sample equals 100. Protein abundance was calculated by summing the intensities of connected peptides. Protein ratios were computed using the geometric median of the protein abundance across biological replicates. Statistical significance of differential protein expressions was assessed using ANOVA, and adjusted p-values were calculated using the Benjamini-Hochberg method. For phosphopeptide analysis, modification site probabilities were calculated using the IMP-ptmRS node in Proteome Discoverer, enabling confident localization of phosphorylation events.

## Acknowledgements

We would like to thank Dr. Justin Mott, Joynob Puspo, and members of the Schott laboratory for their constructive feedback and thoughtful discussions. This work was supported by the National Cancer Institute R21 Grant R21CA279878, the NIAAA R00 Grant R00AA026877, and the NIGMS R35 MIRA Award R35GM150801 (to M.B.S.). Graphical illustrations were created using BioRender.com.

## Conflict of Interest

The authors declare that they have no conflicts of interest.

## Supplementary figure legends

**Supplementary Figure 1.** (A) Fluorescence micrographs of AML-12 cells transfected with TFEB-GFP show rapid nuclear translocation of TFEB following treatment with cAMP agonists (Fsk/IBMX). (B) Quantification of the nuclear-to-cytoplasmic TFEB-GFP ratio reveals a significant increase in nuclear localization within 30 minutes of treatment compared to DMSO control (n = 3; p < 0.05). This data suggests cAMP stimulates TFEB nuclear import in AML-12 cells. (C) Fluorescence micrographs of AML-12 cells transfected with TFEB-GFP and treated with cAMP agonists (Fsk/IBMX), with or without the calcium chelator BAPTA-AM. Chelation of intracellular calcium markedly reduces TFEB nuclear localization within 15 minutes. (D) Quantification of the nuclear intensity of TFEB-GFP confirms a decrease in nuclear TFEB levels in the BAPTA-AM-treated group compared to control. This data suggests cAMP-stimulated calcium promotes TFEB nuclear import in AML-12 cells. (E) Western blot analysis of whole-cell lysates from HeLa cells treated with DMSO (30 min), Fsk/IBMX (30 min), or with or without BAPTA-AM. Blots were probed with a phospho-specific antibody against TFEB S211. (F) Densitometric analysis of panel (E) phospho-S211(normalized to GAPDH) showed reduced S211 phosphorylation in Fsk/IBMX condition and BAPTA-AM but no change following Fsk/IBMX treatment compared to BAPTA-AM condition (n = 3, one-way ANOVA). (G) Western blot analysis of whole-cell lysates from HeLa cells treated with DMSO (30 min), Fsk/IBMX (30 min), or left untreated. Blots were probed with a phospho-specific antibody against TFEB S122. (H) Densitometric analysis of panel (G) phospho-S122 (normalized to GAPDH) showed no change following Fsk/IBMX treatment compared to DMSO control (n = 3, one-way ANOVA).

**Supplementary Figure 2**. (A) Fluorescence micrographs of HeLa cells transiently expressing TFEB–GFP phospho-null mutants (S466A, S467A) or phospho-mimetic mutants (S466E, S467E) treated with Fsk (10 µM) and IBMX (0.5 mM) for 30 minutes or 8 hours. (B) Quantification of nuclear-to-cytoplasmic TFEB–GFP ratios corresponding to panel A. All mutants display robust nuclear retention at 30 minutes compared with baseline (0 hour). By 8 hours, phospho-null (S466A, S467A) and phospho-mimetic (S466E, S467E) mutants show predominant cytoplasmic localization similar to wild-type TFEB, although a modest degree of nuclear retention persists in the phospho-null forms. Data represent mean ± SEM from three independent experiments (n = 3; 100 cells per condition; one-way ANOVA with Tukey’s multiple-comparison test;****P < 0.0001).

**Supplementary Figure 3.** (A) Coomassie blue stain and GFP western blot showed a successful demonstration of GFP-Trap isolation from HeLa cells stably expressing TFEB-GFP versus normal HeLa, where no pulldown artifacts were observed. (B) Heatmap of TFEB–GFP phosphopeptides identified by GFP-Trap pulldown from HeLa cells treated for 30 minutes with DMSO, Fsk (10 µM) + IBMX (0.5 mM), Fsk/IBMX + H89, or H89 alone (50 µM), analyzed by mass spectrometry (n = 3 independent experiments). Phosphorylation at S109, S114, and S122 was significantly increased during the nuclear import phase in a PKA-dependent manner. (C) Fluorescence microscopy of HeLa cells expressing basal TFEB–GFP phospho-null (S109,114,122A) or phospho-mimetic (S109,114,122A) mutants. (D) Quantification of nuclear TFEB levels from panel B shows no significant differences in basal nuclear localization between mutants (n = 3; 100 cells per condition; one-way ANOVA). (E) Representative micrographs of HeLa cells expressing TFEB mutants and treated with DMSO or Fsk + IBMX for 30 min. cAMP stimulation increased nuclear TFEB in both phospho-null and phospho-mimetic mutants. (F) Quantification of nuclear TFEB confirms a significant increase upon Fsk/IBMX treatment compared with DMSO (n = 2; 100 cells per condition). These data indicate that S109, S114, and S122 are not essential for cAMP-mediated TFEB nuclear import.

## Notes

### Competing Interest Statement

The authors have declared no competing interest.

