## Supplementary figures and images for "cAMP promotes acute lysosome biogenesis through TFEB nuclear import-export dynamics"

### Supplementary Figure 1

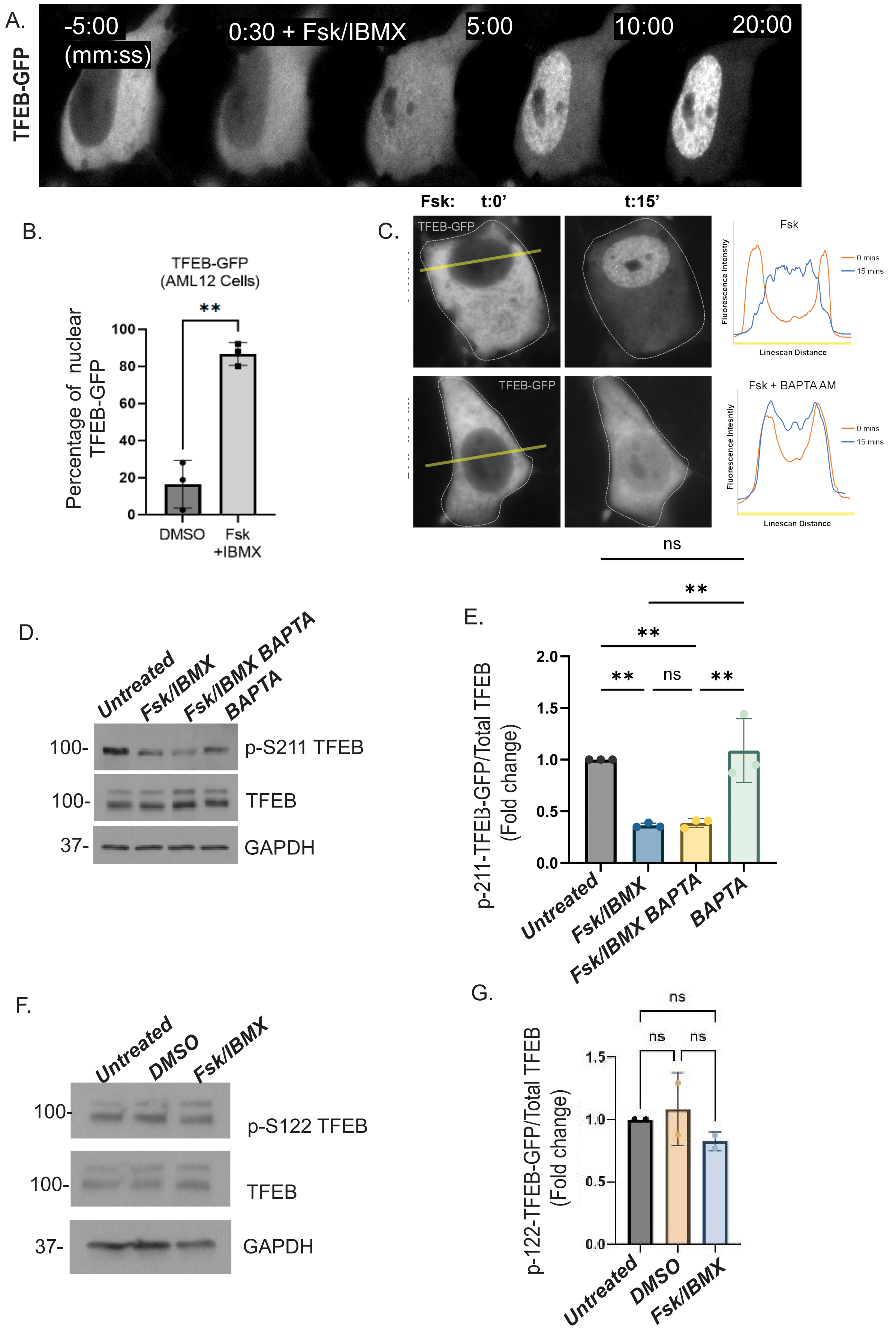

### Supplementary Figure 2

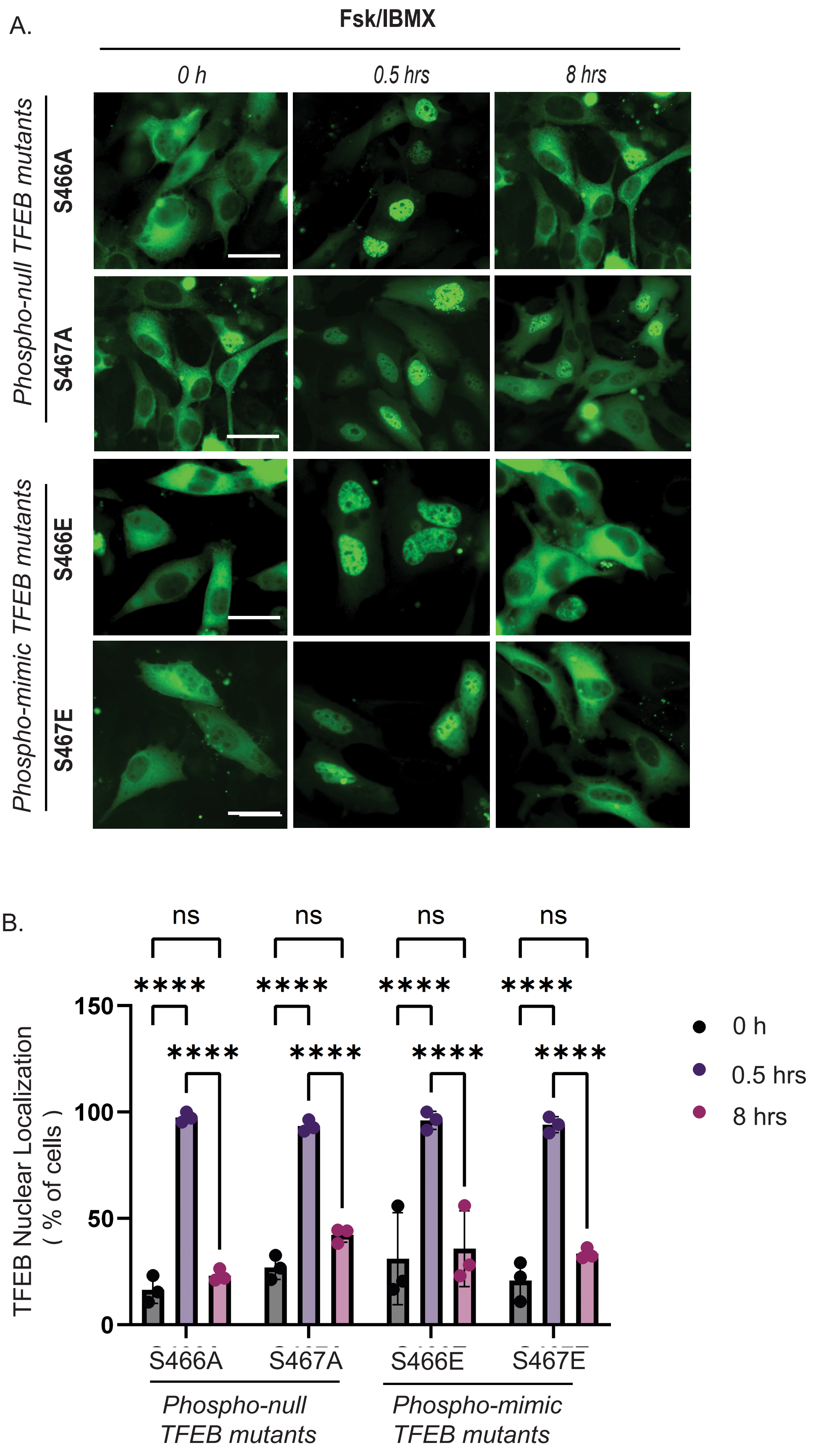

### Supplementary Figure 3

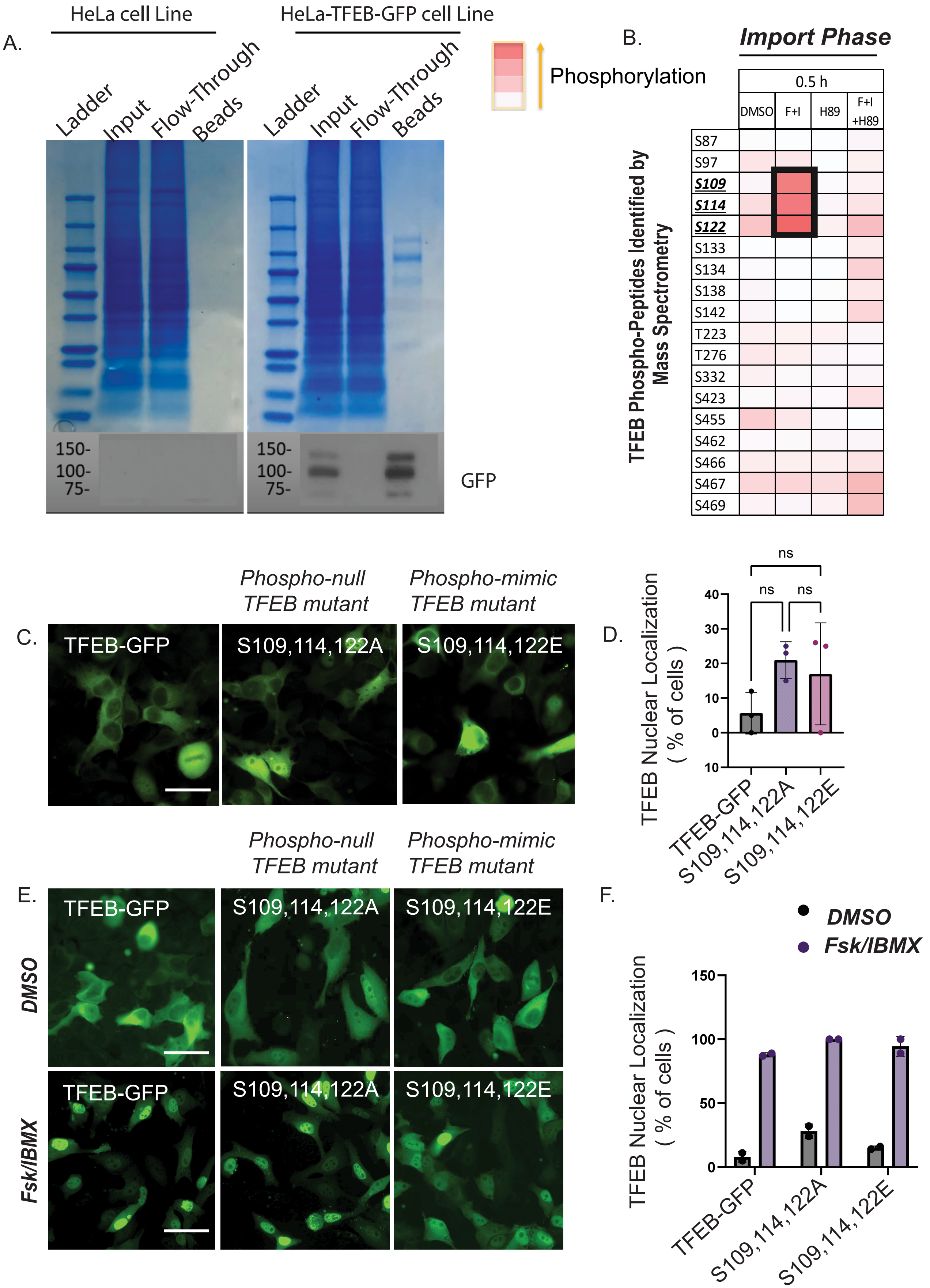
